# Phylogenetic support of *pebS* as a phage-exclusive auxiliary metabolic gene

**DOI:** 10.64898/2026.01.23.701310

**Authors:** Nina Baeuerle, Nicole Frankenberg-Dinkel, Anne Kupczok

## Abstract

Marine picocyanobacteria, including the genera *Prochlorococcus* and *Synechococcus*, are major contributors to oceanic photosynthesis and global primary production. Their populations are influenced by T4-like cyanophages, which frequently encode auxiliary metabolic genes (AMGs) capable of altering host metabolism during infection. One such AMG, *pebS*, encodes a ferredoxin-dependent bilin reductase (FDBR) phycoerythrobilin (PEB) synthase, which converts biliverdin IXα to PEB. In contrast, cyanobacteria perform a two-step reaction using the FDBR enzymes PebA (15,16-dihydrobiliverdin:ferredoxin oxidoreductase) and PebB (PEB:ferredoxin oxidoreductase), whereas *pebS* has not been reported in cyanobacterial genomes. Here, we re-evaluated whether *pebS* is truly restricted to cyanophages by searching the Ocean Gene Atlas and all available cyanobacterial genomes at NCBI using a cyanophage-derived PebS sequence as query. Using protein phylogenies, we find that most search hits group with PebA or PebB, while few sequences from cyanobacterial genome assemblies were confirmed to belong to PebS based on phylogenetic placement. However, genomic context analysis of these *pebS* sequences revealed that they are phage-derived, consistent with cyanophage infection at the time of sampling. In conclusion, our results support that *pebS* is absent in cyanobacterial genomes, raising questions about the evolutionary and biochemical rationale for the two-step reduction of biliverdin IXα to PEB in these organisms.

## Introduction

The marine picocyanobacteria *Prochlorococcus* and *Synechococcus* are the most abundant photosynthetic organisms in the marine environment (Flombaum *et al*. 2013; Scanlan and West 2002) and thereby play an important role in primary production in the oceans. The populations of *Prochlorococcus* and *Synechococcus*, as well as other marine cyanobacteria, are affected by the infection with cyanophages (Dekel-Bird *et al*. 2015; Warwick-Dugdale *et al*. 2019). Many cyanophages are T4-like phages possessing genomes that, in addition to the conserved T4 core genes, encode diverse auxiliary metabolic genes (AMGs). Many AMGs are host-like and are thought to have been acquired by the phage via horizontal gene transfer from their host (Bryan *et al*. 2008; Zeng and Chisholm 2012). These genes are hypothesized to modulate host metabolism during infection, thus conferring an advantage in the production of phage particles (Puxty *et al*. 2015; Hurwitz and U’Ren 2016). Cyanophage AMGs frequently encode proteins involved in photosynthesis, with *psbA*, encoding the D1 subunit of Photosystem II, being the most abundant one (Lindell *et al*. 2005; Sullivan *et al*. 2006). In addition, genes encoding proteins involved in the biosynthesis of phycobilins are frequently found. These linear tetrapyrrole pigments are essential for the cyanobacterial light-harvesting complexes, the phycobilisomes (PBS) (Dammeyer and Frankenberg-Dinkel 2008). Among them are the genes heme oxygenase (*ho1)* and phycocyanobilin:ferredoxin oxidoreductase (*pcyA)*, both encoding enzymes with the same activity as their host counterparts (Sullivan *et al*. 2005; Dammeyer *et al*. 2008). The heme oxygenase is involved in the cleavage of the heme macrocycle to the first linear tetrapyrrole biliverdin IX α (BV), and PcyA, a ferredoxin-dependent bilin reductase (FDBR) reducing BV to the light harvesting chromophore phycocyanobilin (Dammeyer *et al*. 2008; Frankenberg and Lagarias 2003). In the first sequenced cyanophage genomes, an additional AMG encoding a putative FDBR was identified (Sullivan *et al*. 2006). Interestingly, the FDBR PebS, found in certain *Prochlorococcus* and *Synechococcus* infecting phages, catalyzes a reaction that requires two separate enzymes in the host (Dammeyer and Frankenberg-Dinkel 2008). Cyanobacteria perform two subsequent two-electron reductions by the FDBRs 15,16-dihydrobilinverdin (DHBV):ferredoxin oxidoreductase PebA and phycoerythrobilin (PEB):ferredoxin oxidoreductase PebB. The FDBR PebA reduces the substrate BV at the 15,16-methine bridge of the tetrapyrrole to the intermediate 15,16-DHBV which is directly converted to PEB by PebB (Dammeyer and Frankenberg-Dinkel 2006). As demonstrated *in vitro*, PebS catalyzes the complete four-electron reduction of BV to PEB in a single, one-step reaction. Owing to its high amino acid sequence similarity to PebA, PebS was originally misannotated as PebA (Dammeyer *et al*. 2008). However, unlike HO1 and PcyA, PebS does not mirror the reaction performed by its host, raising the question on its origin and evolution. In line with this, no cyanobacteria have been identified to possess the *pebS* gene. Here, based on publicly available metagenomes and cyanobacteria assemblies, we re-evaluate the genomic distribution of *pebS* and show that it is exclusively encoded by phages.

## Methods

### Preparing a dataset of PebA and PebB sequences

For better comparability of PebA, PebB, and PebS, several PebA and PebB amino acid sequences were retrieved from the UniProt database (hereinafter referred to as test data set, Table S1). Five sequences from *Prochlorococcus* and *Synechococcus* were used for PebA and PebB, respectively. Five sequences were used for PebS.

### Preparing an HMM from PebS sequences

Amino acid sequences of the FDBR PebS were obtained by searching in the ClusteredNR database (date 16 September 2025) using the blastp webservice at NCBI (Altschul *et al*. 1990) with the PebS sequence of the cyanophage P-SSM2 (GenBank accession: ACY75934.1) as input. The word size was reduced to 2, other parameters were left at default. Hits annotated as PebS belonging to cyanophages were downloaded and aligned with the original PebS using the web service MAFFT (Kuraku *et al*. 2013; Katoh, Rozewicki and Yamada 2019) v7.511 with default parameters. Additionally, for a more sensitive search, a hidden Markov model (HMM) was created from the alignment using hmmbuild rev3.1b2 (Finn, Clements and Eddy 2011; Hoang *et al*. 2018).

### Screening of PebS in the Ocean Gene Atlas (OGA)

The HMM was used for screening for bacterial PebS sequences in metagenome assemblies from the ocean gene atlas v2.0 (Vernette *et al*. 2022) (dataset OM-RGCv2 +G) with default parameters. The resulting bacterial sequences were extracted and aligned as described above. A phylogenetic tree was constructed from the alignment with IQ-TREE v3.0.1 (Hoang *et al*. 2018; Wong *et al*. 2025) (-B 1000) and the best selected model.

### Search for pebS in cyanobacterial genomes (NCBI database)

Sequences of cyanobacteria (Cyanobacteriota (blue-green bacteria)) were obtained from the National Center for Biotechnology Information (NCBI) genome database (September 24, 2025). All genomes at all assembly levels were used. The completeness of the genomes was determined using checkm2 v1.1.0 (Chklovski *et al*. 2023). The contigs of the individual genomes were tested for possible phage infection by detecting phage genomes with GeNomad v1.9.0 (Chen *et al*. 2024).

The encoded protein sequences were used to create a cyanobacterial database with blast v2.16.0+. The database was then screened for PebS sequences using the cyanophage P-SSM2 PebS as input sequence. The word size was therefore reduced to 2, other parameters were left at default. The resulting sequences were aligned using the web service MAFFT v7.511 with the FFT-NS-i method. A phylogenetic tree was constructed from the alignment with IQ-TREE v3.0.1 (-B 1000) and the best selected model.

### Search for pebA and pebB in cyanobacterial genomes

To check whether *pebA* and *pebB* sequences are also present in the genome alongside *pebS*, the genomes in which *pebS* sequences were found were also screened for pebA and *pebB*. For this purpose, the PebA and PebB sequences from *Synechococcus* sp RS9916 were blasted against the translated genomes (webservice tblastn, at NCBI).

## Results

### PebA, PebB and PebS form distinct groups in a phylogenetic tree

To check cyanobacterial genomes for the presence of *pebS*, we first aim to establish the relationships between the FDBRs PebS, PebA, and PebB. To this end, a test data set with known amino acid sequences of cyanobacterial PebA and PebB, as well as annotated PebS sequences from cyanophages was used (Table S1). The phylogeny constructed from these sequences supports that PebA, PebB, and PebS form distinct groups (Figure S1, each with bootstrap support at least 95), confirming previous results (Dammeyer *et al*. 2008; Ledermann, Béjà and Frankenberg-Dinkel 2016). Since these FDBRs show high sequence similarities, they can be found simultaneously in a blastp search and thus a phylogenetic placement of newly detected FDBRs will be used to distinguish them.

### PebS sequences are absent in bacterial assemblies from the Ocean Gene Atlas

To investigate whether *pebS* genes can also be found in cyanobacterial genomes, several databases were systematically screened. In a first approach, the Ocean Gene Atlas (Tara Ocean) (Vernette *et al*. 2022; Katoh, Rozewicki and Yamada 2019) was searched. For this purpose, a Hidden Markov Model (HMM) constructed from 30 cyanophage PebS amino acid sequences was used as input. The search resulted in 1,038 amino acid sequences, of which 95 sequences were classified as belonging to cyanobacteria. However, after multiple sequence alignment (MSA) and the construction of a phylogenetic tree, it became apparent that none of the sequences are actual PebS (Fig. S2). Instead, the search found sequences from the related FDBRs PebB (13 sequences) and PebA (79 sequences). In addition, the phylogenetic tree contains an additional group of three sequences that cannot be clearly assigned to any of the three FDBRs.

### Five pebS sequences can be found in cyanobacterial assemblies from NCBI

In a second approach, a database was created using all cyanobacterial genome sequences available at NCBI. All assembly levels were included, resulting in a database with the protein sequences of 10,892 cyanobacterial genomes (Table S2). Subsequently, a search against this database was performed with blastp using PebS from the cyanophage P-SSM2 as the input sequence. A phylogenetic tree was constructed from the top 500 resulting sequences aligned with the test data (Figure S2). Based on the phylogeny, two main groups can be identified; a large cluster corresponding to PebA (495 sequences excluding the test dataset) and a small cluster corresponding to PebS (five sequences excluding the test dataset). Note that the PebS sequences were initially found as the top 5 hits of the blast search, having a lower e-value than PebA sequences, thus, we conclude that we found all PebS in cyanobacteria. The phylogeny of the test data with the five identified PebS sequences (Fig. 1) also supports the different group and the five sequences belong to PebS.

**Figure 1:**
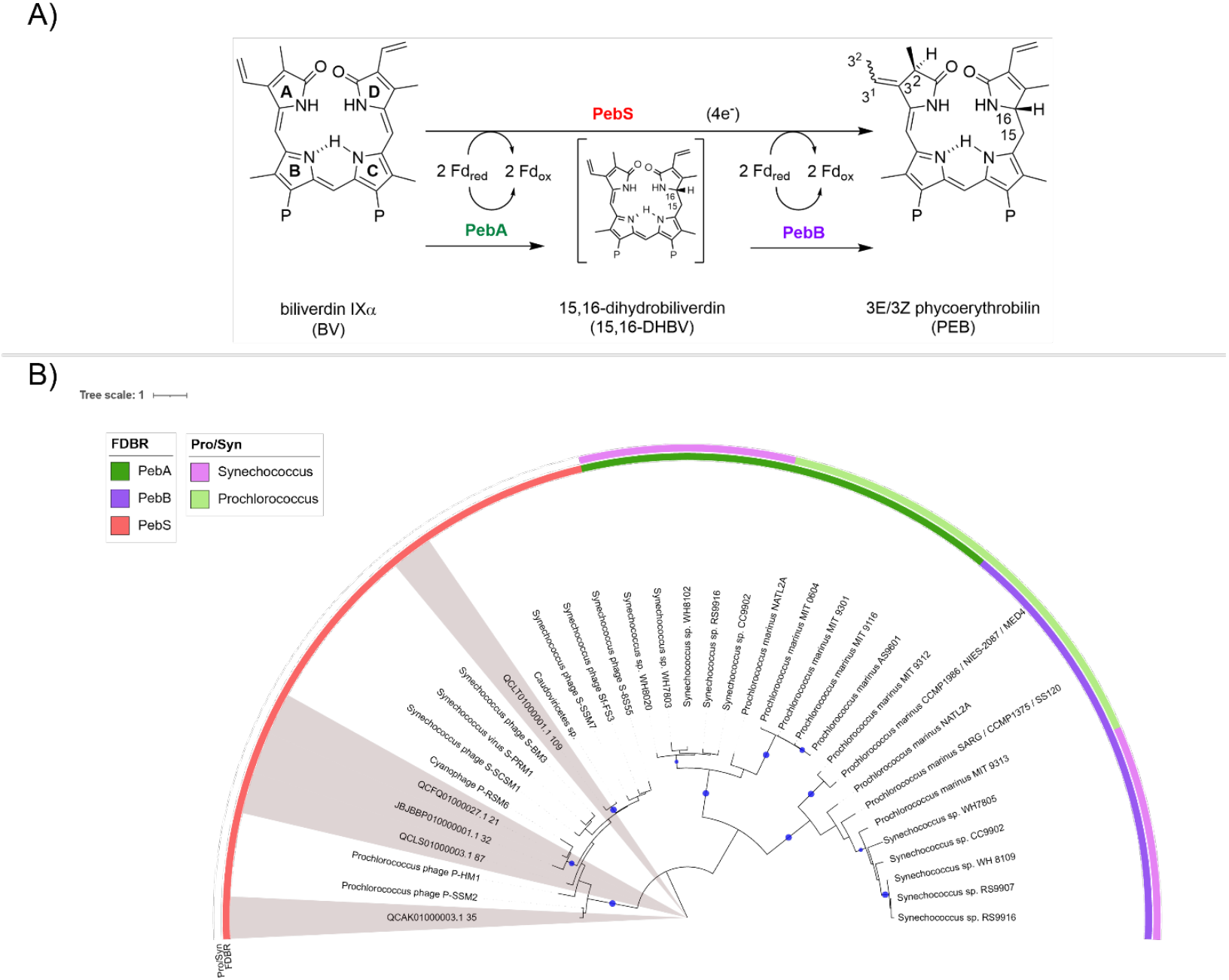
PEB synthesis and phylogenetic tree of PebA, PebB and PebS. **A)** Reduction of biliverdin via the intermediate DHBV by the enzymes PebA and PebB encoded in cyanobacteria; one-step reduction via the cyanophage-encoded enzyme PebS. **B)** Midpoint rooted maximum likelihood phylogenetic tree with PebS data from NCBI (red) sequences and a test dataset consisting of PebA (dark green) and PebB (purple) sequences from *Prochlorococcus* (pink) and *Synechococcus* (light green). PebS sequences obtained from the cyanobacterial genome sequence search are marked with a grey background. The tree was constructed using IQ-Tree and visualized with iTOL as midpoint rooted (Letunic and Bork 2011, 2024). Bootstrap support ≥ 95 is indicated with a blue circle. The scale bar indicates the average number of amino acid substitutions per site.

The five newly identified PebS sequences are all from the marine environment and originate from one *Synechococcus* and four *Prochlorococcus* genomes, respectively (Table 1). The *Synechococcus* genome is a metagenome assembled genome (MAG) with a completeness estimated with checkm2 of 100%. The four *Prochlorococcus* genomes are single-cell amplified genomes (SAGs) with completeness ranging between 12 % and 96%. The overall contamination rate of the genomes is low, at a maximum of 1.5%.

**Table 1:**
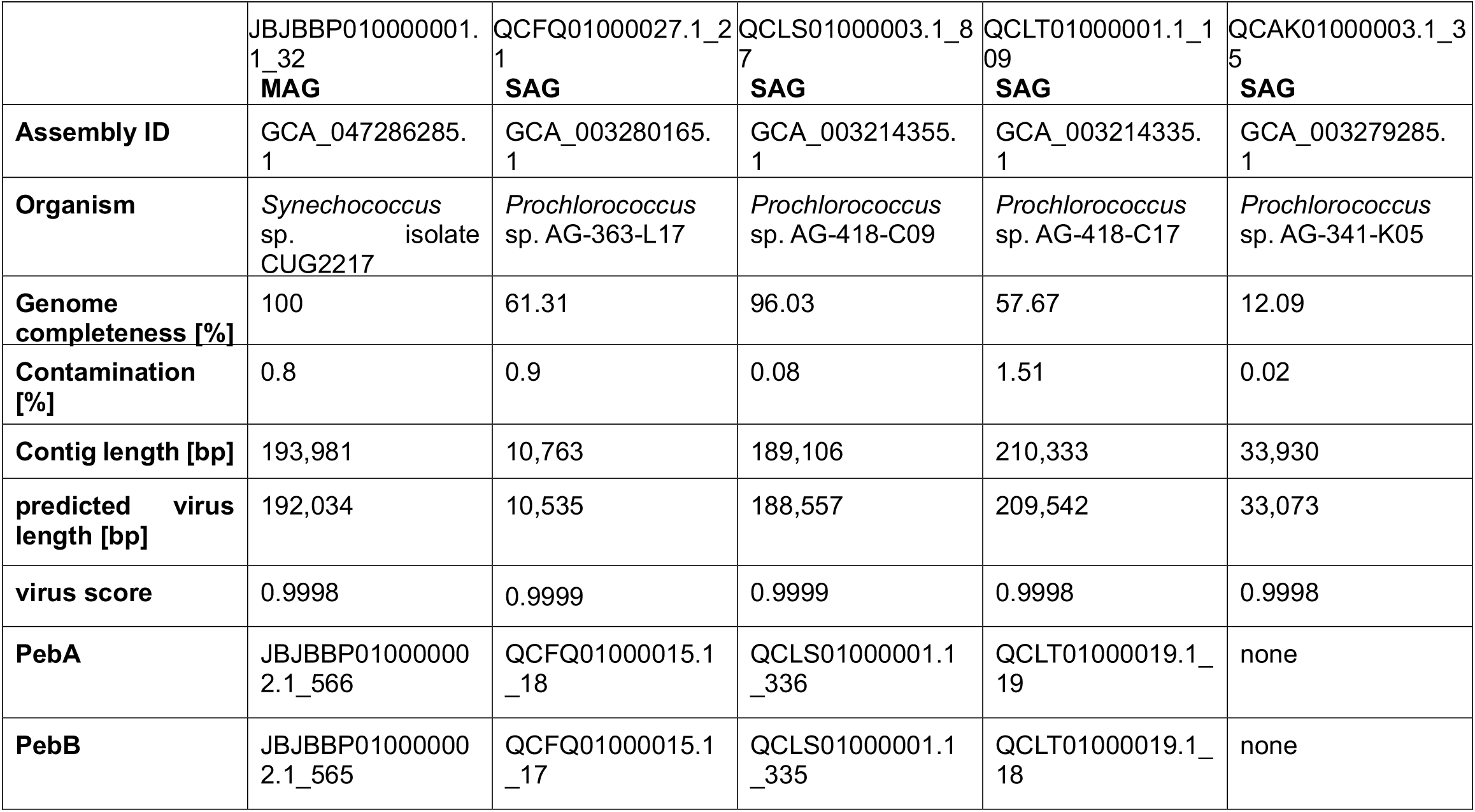
PebS sequences detected in cyanobacterial genomes. For each of the sequence numbers, the assembly ID as well as the genome type is given. MAG - metagenome assembled genome; SAG – single-cell amplified genome. The genome completeness as well as the contamination was obtained using checkm2. The predicted virus length as well as the virus score was obtained using geNomad. The genomes were additionally checked for PebA and PebB sequences.

*All pebS sequences detected in cyanophage genomes can be traced back to phage DNA* Since *pebS* was so far only known in cyanophage genomes, we next wanted to test whether the detected *pebS* sequences are embedded within phage genomes that might have been included in the sequencing during an active phage infection. To this end, the genomes were additionally examined using geNomad to detect phage DNA. The geNomad results show that all contigs containing the *pebS* sequences are contigs of phage origin (Table 1). In all cases, the predicted virus length fills nearly the entire contig and the virus score is close to one, clearly identifying them as phages. In addition to the *pebS* sequence, *pebA* and *pebB* were also found in the genomes in four of five cases. For the genome where *pebA* and *pebB* could not be found, the genome completeness is very low (12%); thus, no statement can be made about the presence or absence of *pebA* and *pebB* in the genome.

## Discussion

Here, we set out for searching the phage AMG *pebS* in bacterial genomes. First, we searched in the ocean gene atlas, where we could not find it. Second, among 10,892 publicly available cyanobacteria genome assemblies in NCBI, we detected five p*ebS* sequences that we confirmed to encode PebS due to their phylogenetic placement. However, a closer examination of these genome assemblies revealed that they originated from single-cell genomics or metagenomics of environmental samples. Furthermore, the p*ebS* sequences can all be found on phage contigs; thus, the samples likely included phage DNA. Since no temperate cyanophages are known to exist (Sullivan, Waterbury and Chisholm 2003) and since the predicted virus length covers nearly the complete contigs with *pebS*, the detected phage genomes likely originate from a lytic phage infection. The high prevalence of lytic phage infections within public bacterial assemblies was recently discovered and challenges classical phage lifestyle categories (Perfilyev *et al*. 2026). Here we show how the presence of lytic phage genomes in bacterial assemblies also complicates the evolutionary analysis of AMGs.

From the presented analysis, we can conclude that no known cyanobacteria genomes have the *pebS* gene. Instead, they harbor *pebA* and *pebB*. Nevertheless, we cannot rule out the possibility that cyanobacteria with *pebS* exist and have simply not yet been found. Further possibilities for searching for PebS in currently known sequence data could be explored in the future. For example, metagenome assemblies could be searched with Logan (Chikhi *et al*. 2025) using diverse *pebS* gene sequences.

Our findings raise the question how the AMG *pebS* evolved. PebS shows a strong similarity to PebA. In addition, the mutation of a crucial aspartate amino acid can alter the function of PebS so that it only performs the reduction of BV to 15,16-DHBV (Busch *et al*. 2011). This suggests that *pebA* could have evolved from *pebS* through mutation, or vice versa. Thus, one possibility is that phages have acquired *pebA* from their cyanobacterial host via horizontal gene transfer, which then evolved into *pebS*. However, it is also conceivable that cyanobacteria originally possessed *pebS*, which was acquired by the phage. After the horizontal gene transfer, gene duplication in the cyanobacterial host could have resulted in the two genes *pebA* and *pebB*. This scenario is also supported by the fact that *pebA* and *pebB* genes occur together in a bicistronic operon, the *pebAB*-operon (Dammeyer and Frankenberg-Dinkel 2006; Aras *et al*. 2020). Neither evolutionary scenario has been proven conclusively and although extensive phylogenetic analysis has been performed on the whole FDBR family in order to investigate their origin, there is still lacking evidence for the origin of PebS (Rockwell, Martin and Lagarias 2023).

Regardless of these two possibilities, another question is why it is advantageous for cyanophages to carry the one-step enzyme while cyanobacteria split the reaction into two enzymatic steps. The reason for this could be that it is advantageous for viruses to keep their genome as small as possible, since a smaller genome allows faster genome replication and thus faster release of new virus particles (DiMaio 2012; Hatfull and Hendrix 2011). Indeed, the gene *pebS* only needs half of the genetic material compared to *pebA* and *pebB*. Furthermore, why is it advantages for cyanobacteria to split the reaction into two enzymatic steps? One explanation for this would be a tighter regulation of PEB production. Another possibility would be the potential use of the reduction product of BV IXα (Beale and Cornejo 1991; Wedemayer *et al*. 1992). However, no use of 15,16-DHBV in cyanobacteria has been demonstrated to date. This contrasts with findings in cryptophytes, where it has been shown that 15,16-DHBV binds to their specific phycobiliproteins (Wedemayer *et al*. 1992). Another reason for dividing the reaction into two steps in cyanobacteria could be the possibility of tighter regulation through, for example, feedback inhibition (Dammeyer *et al*. 2008). Thus, despite the availability of a single-enzyme alternative, cyanobacteria rely exclusively on the PebA-PebB pathway for PEB biosynthesis, and the reasons for the absence of *pebS* from their genomes remain to be elucidated.

## Acknowledgements

This work was funded within the DFG priority programme SPP2330 to NFD.

## Abbreviations

AMG: auxiliary metabolic gene
BV: biliverdin
DHBV: dihydrobiliverdin
HMM: hidden markov model
MAG: metagenome assembled genome
MSA: multiple sequence alignment
NCBI: National Center for Biotechnology Information
PBS: phycobilisome
PEB: phycoerythrobilin
SAG: single amplified genome

## Notes

### Competing Interest Statement

The authors have declared no competing interest.

