## Supplementary material for "Phylogenetic support of *pebS* as a phage-exclusive auxiliary metabolic gene": Figure S

#### Supplementary figures

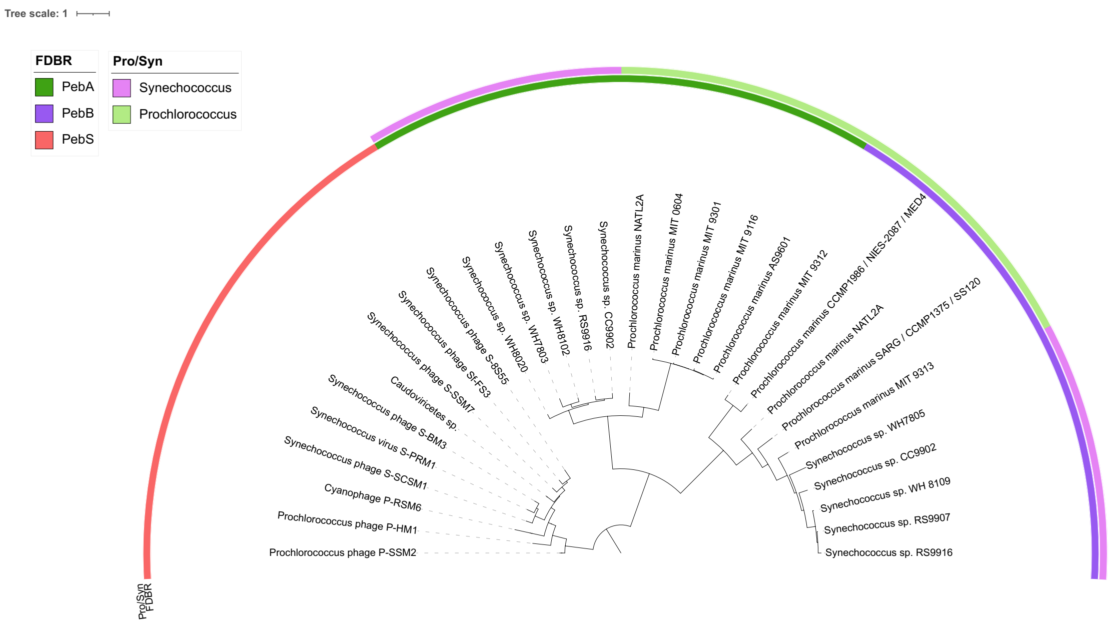


**Figure S1: Midpoint rooted maximum likelihood phylogenetic tree with PebS (red), PebB (purple) and PebA (green), test dataset. The** tree was constructed using IQ-Tree and visualized with iTOL.

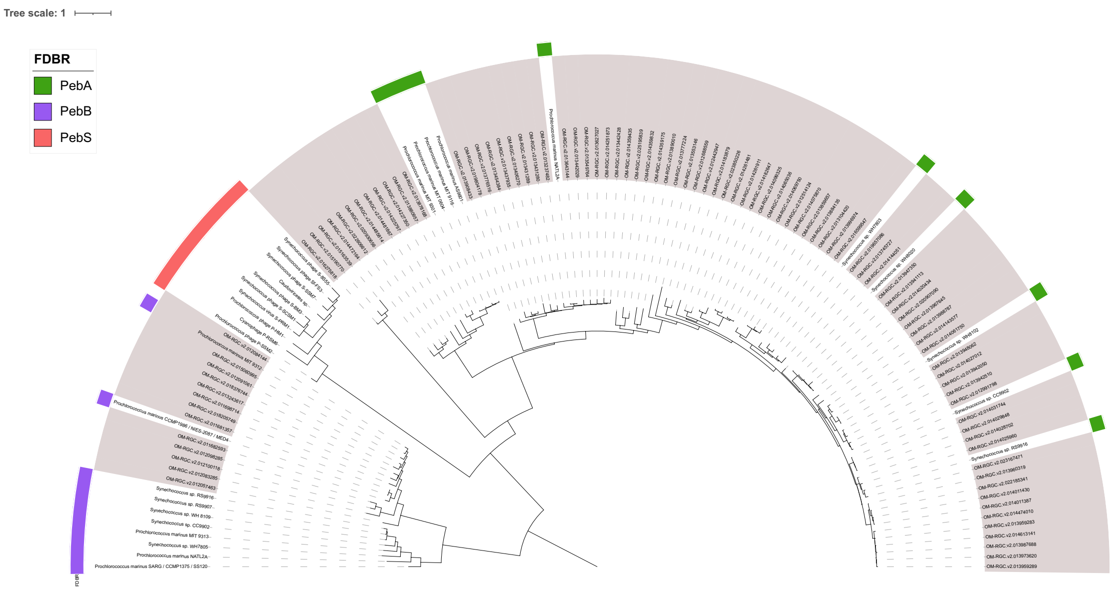


**Figure S2: Midpoint rooted maximum likelihood phylogenetic tree with PebS (red), PebB (purple) and PebA (green) data from the OGA sequences (grey background) and a test dataset (marked with the respective colour). The** tree was constructed using IQ-Tree and visualized with iTOL.

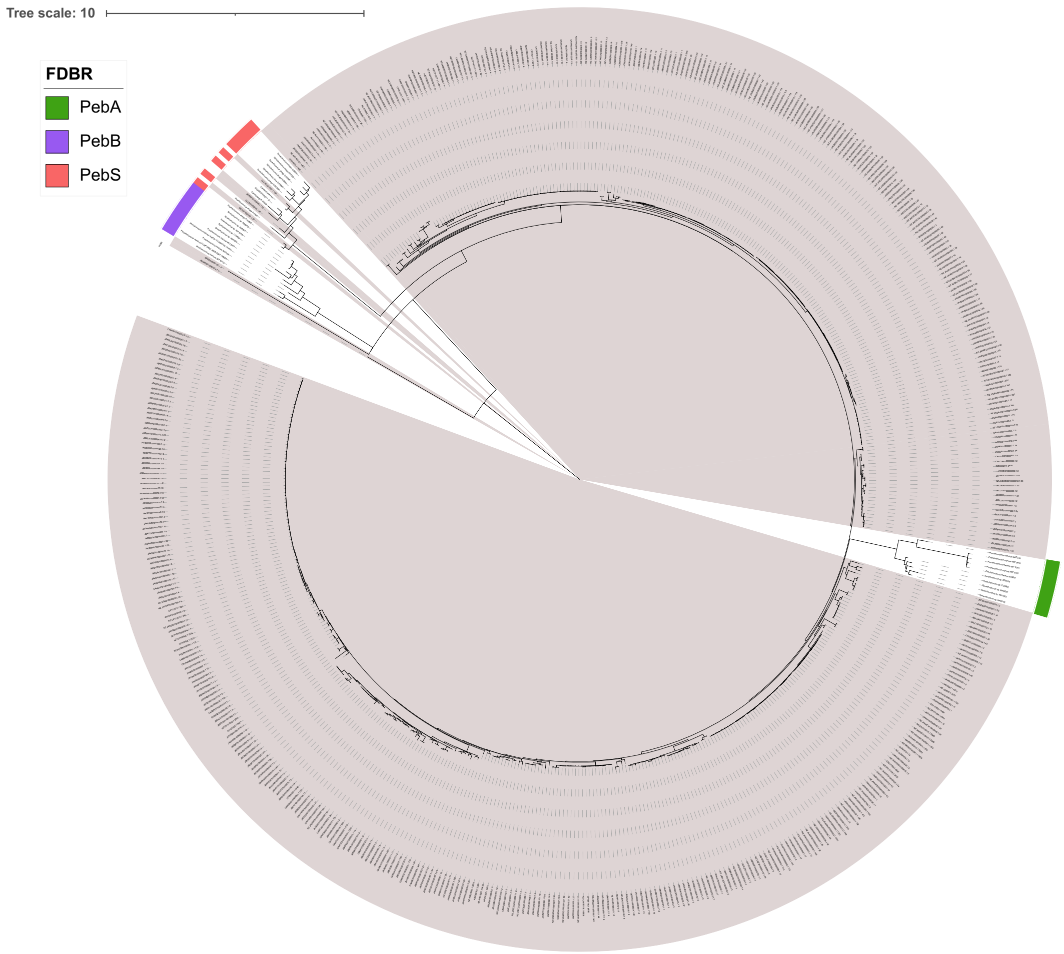


**Figure S3: Midpoint rooted maximum likelihood phylogenetic tree with PebS (red), PebB (purple) and PebA (green) data from NCBI sequences (grey background) and a test dataset (marked with the respective colour). The** tree was constructed using IQ-Tree and visualized with iTOL.

### Supplementary Tables

**Table S1: Test dataset for PebA, PebB and PebS with accession number.**

**Table S2: Genomes used for the analysis of pebS in cyanobacterial genomes from the NCBI**
